# Opportunity or catastrophe? Effect of sea salt on host-parasite survival and reproduction

**DOI:** 10.1101/2021.06.03.446887

**Authors:** Ao Yu, J. Trevor Vannatta, Stephanie O. Gutierrez, Dennis J. Minchella

**Author notes:** (AY); (AY), (JTV); (JTV). **Author contributions** AY, JTV, SOG, and DJM conceived and designed the study. AY, JTV, and SOG conducted the study. AY and JTV analyzed the data. AY, JTV, SOG, and DJM contributed to writing and revising the manuscript. **Data availability:** All data and code used for this study are available at https://github.com/vanna006/sea-salt-schisto.

## Abstract

Seawater intrusion caused by anthropogenic climate change may affect freshwater species and their parasites. While brackish water certainly impacts freshwater systems globally, its impact on disease transmission is largely unknown. This study examined the effect of artificial seawater on host-parasite interactions using a freshwater snail host, *Biomphalaria alexandrina*, and the human trematode parasite *Schistosoma mansoni*. Four components were analyzed to evaluate the impact of increasing salinity on disease transmission: snail survival, snail reproduction, infection prevalence, and the survival of the parasite infective stage (cercariae). We found a decrease in snail survival, snail egg mass production, and snail infection prevalence as salinity increases. However, cercarial survival peaked at an intermediate salinity value. Our results suggest that seawater intrusion into freshwaters has the potential to decrease schistosome transmission to humans.

**Author Summary:** Climate change will have numerous impacts on many systems, including host-parasite systems. One mechanism by which climate change with impact host-parasite interactions is by rising sea levels flooding coastal regions, increasing salinity in many freshwaters. Host-parasite interactions are a key component of freshwater ecosystems, but the effects of sea water intrusion on host-parasite dynamics are largely unknown. In this study, we quantify the effects of sea salt concentration on the model host-parasite system, *Biomphalaria alexandrina* and *Schistosoma mansoni*. We demonstrate a significant, negative relationship between sea salt concentration and host-parasite survival and reproduction. The increase in freshwater salinity associated with sea level rise has the potential to decrease parasite transmission and disease burden in humans and wildlife.

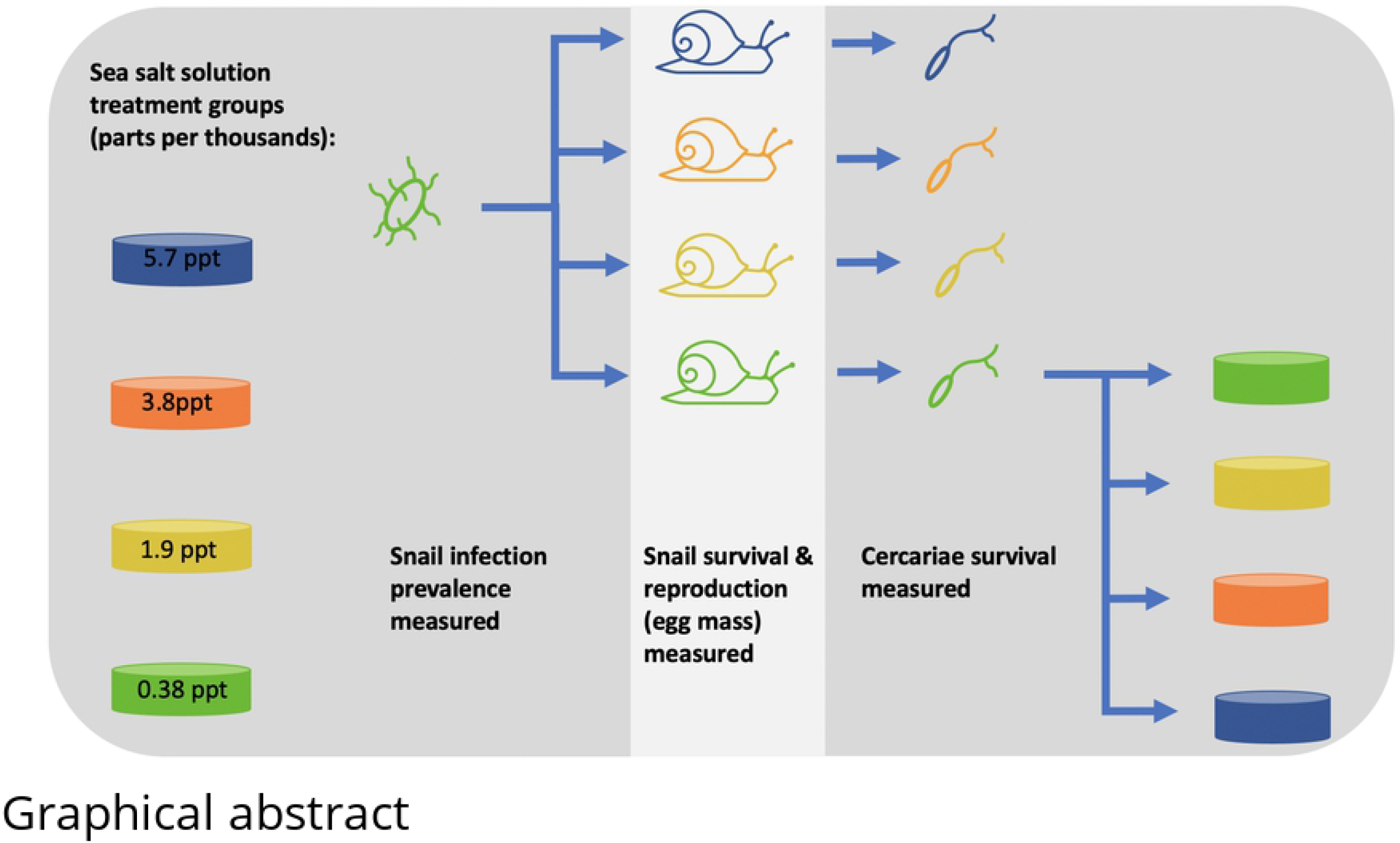

## Introduction

Anthropogenic climate change, associated with increasing greenhouse gases, can change many environmental factors including temperature, pH, precipitation, as well as salinity in both terrestrial and aquatic systems. Increasing salinity in freshwater systems is often caused by rising sea levels and seawater intrusion [1,2]. Salt concentration (salinity) is a crucial abiotic factor that impacts many aspects of biotic interactions [3]. Increases in salinity of freshwaters will undoubtedly result in changes in organismal growth, reproduction, and survival, impacting entire food webs, and thus host-parasite interactions.

Parasites play a key role in freshwater communities [4], and they are commonly recognized for their ability to modify the growth, reproduction, and survival of their hosts [5]. They are also the etiologic agents of human disease. Therefore, understanding how abiotic factors and parasitic diseases interact in the context of climate change, more specifically increasing salinity, is critical in order to understand human disease transmission.

Rising seawater levels and seawater intrusion are affecting numerous bodies of water including the Nile River Delta. The salinization of coastal land in the Nile Delta is caused by the decrease in the Nile River’s freshwater levels due to human activity and the increasing sea levels in the Mediterranean Sea [6]. Seawater will submerge large areas in the coastal zone of the Nile Delta in the near future, impacting host-parasite interactions in the region [7].One parasitic disease of humans acquired in the Nile Delta of Egypt is schistosomiasis, which is caused by the trematode *Schistosoma mansoni* [8]. Approximately, 12.7 million infected people are clustered in the Middle East and North Africa region, and Egypt’s share of the burden is about 7.2 million [9]. Parasite transmission occurs when humans come in contact with secondary parasite larvae, called cercariae, which are released from infected snails in freshwaters. In areas with perennial irrigation such as the Nile Delta and the Nile River Valley, schistosomiasis prevalence is high (60% infection rate). In contrast, the infection rate is relatively low (6%) in districts of basin irrigation (commonly known as annual flooding) [10]. Development and shift from basin irrigation to perennial irrigation in Egypt has resulted in year-round availability of water in many districts making the likelihood of infection high [11], but the impact of seawater intrusion on parasite transmission in this system is unclear.

This laboratory study explores the effect of increasing salinity (as artificial seawater) on host and parasite success using the Egyptian strain of *Schistosoma mansoni* and its snail intermediate host, *Biomphalaria alexandrina*, to simulate the conditions in the Nile Delta of Egypt. In order to assess the success of parasite transmission, we exposed snail hosts to the trematode parasite and monitored snail survival, snail egg mass production, infection prevalence, and parasite (cercarial) survival. We predicted lower snail survival and egg mass production as snails become stressed by osmoregulation and allocate less energy to reproduction [12,13]. We further expected to observe lower parasite infection prevalence in snails as salt concentration increased, since the parasite larvae that infect snails (miracidia) may be less successful due to increased energy expenditures required maintain osmolarity [14,15]. Finally, we predicted lower cercarial (the parasite larvae which infects humans) survival as salt concentration increased due to increases in energy expenditures on osmoregulation [16] and fewer available resources from stressed hosts [17].

## Materials and Methods

### Experimental reagents

Various sea salt brands were assessed to determine which most closely resembled the mineral concentrations of seawater (supplement **Figure S1**; **Table S1**). Based on these comparisons, Instant Ocean^®^ Sea Salt was used to replicate seawater.

Salinity of the Mediterranean Sea is approximately 38 parts per thousand (ppt) [18], which we considered saturated seawater (100% salt solution) in this experiment. Salinity treatment groups consisted of 1% (0.38 ppt), 5% (1.9 ppt), 10% (3.8 ppt), and 15% (5.7 ppt) seawater solutions, which were chosen based on pilot data suggesting seawater concentrations of 20% and higher rapidly killed the snail hosts. The four salinity treatments were prepared from 100% stock solution (38 ppt) diluted with well water (the 1% solution), and the salinity of the final solutions were verified by using a handheld refractometer (Premium Aquatics). The 1% (0.38 ppt) salinity treatment was designated as the control for this experiment as this was the salinity of well water used for dilutions.

### Experimental strain maintenance

We used *Biomphalaria alexandrina* snails and *Schistosoma mansoni* parasites, which both originated from Egypt. *B. alexandrina snails* were raised under controlled laboratory conditions (∼25 °C, 12 hour light/12 hour dark). The *S. mansoni* life cycle was maintained for the experiment using *B. alexandrina* snails and male Balb/c mice [19].

In order to limit the shock induced by sudden seawater exposure, 60 snails were acclimated for five days in each salinity treatment (total of 240 snails) before inclusion in this study and subsequent *Schistosoma* infections. In a pilot study, five days was shown to sufficiently control for initial mortality. During the experiment, snails were housed in individual 225 mL jars with Styrofoam (for egg laying) and were fed romaine lettuce *ad libitum*. Water in each jar was changed weekly to maintain salinity levels. Egg masses were counted weekly and removed from jars during cleaning.

### Snail infections

Mice were euthanized in accordance with Purdue Animal Care and Use Committee Protocol # 1111000225 to isolate *Schistosoma* eggs. The collected eggs were placed in freshwater for approximately 60 minutes to allow the miracidia (the infective stage for the snails) to hatch. The resulting miracidia were transferred to well plates containing snails and 10 mL of their respective salinity treatment. Each snail was exposed individually using eight miracidia in the corresponding salinity treatment and left overnight.

### Data collection

Snail survival and egg masses laid were recorded weekly for eight weeks. In weeks four to seven post parasite exposure, snails from each group were transferred into well plates filled with 10 mL of corresponding salinity treatment solution and cercarial release was observed. After one hour under artificial light, 1 mL of mixed solution was drawn from each well, and the number of cercariae was counted for each snail [20]. Snails in which at least 10 cercariae were detected from the randomized 1 mL out of 10 mL were designated as infected. Only 13 out of 948 observations (1.37%) resulted in the snail being assigned as uninfected while having countable cercariae from the randomized 1 mL sample. Thus, cross-contamination was infrequent, and likely had little impact on our results.

Eight weeks after parasite exposure, cercariae from the 0.38 ppt salinity (control) treatment were collected and were used to examine cercarial survival in various salt solutions. Cercarial survival was assessed using only cercariae from the 0.38 ppt salinity treatment due to high snail mortality and low infection prevalence in the other treatment groups. Approximately 20 cercariae were placed into individual wells with 1ml of the four salinity treatment solutions. Cercariae survival was checked at 4h, 8h, 12h, and 24h after release from the snail, and cercariae were removed if little to no movement was detected. Following the 24h check, all cercariae were euthanized with ethanol and counted to determine the exact number of cercariae in each trial.

### Statistical analyses

Snail survival was analyzed with a proportional hazard regression model using the survminer and survival packages in R [21,22]. Snail egg mass production was only compared between uninfected snails to remove the confounding influence of parasitic castration. Even after removal of infected snails, the data had considerable zero-inflation. To account for this, we used a zero-inflated, negative binomial, mixed effects model created in the glmmTMB package to analyze snail reproduction [23]. Differences in infection prevalence between salinity treatments were examined using a binomial General Linear Model. Cercarial survival was analyzed with the coxme package using a mixed effects proportional hazard model[24]. This was done to control for non-independence as multiple cercariae were taken from the same snail. In these models, salinity treatment was used as a fixed factor and individual snail ID was used as a random factor. All pairwise comparisons were made using the emmeans package with the multivariate t distribution for p value correction[25]. All analyses were performed in R version 3.6.3 [26].

## Results

### Effect of seawater on snail survival

Survival probability of infected and uninfected snails in the 0.38 ppt control treatment did not differ significantly over the 8-week experiment (P > 0.05). Therefore, we compared snail survival probability across all salinity treatments regardless of infection status (**Figure 1**). All salinity treatments were significantly different from each other except for 3.8 ppt and 5.7 ppt (P > 0.05). Snails in the 0.38 ppt control treatment had the highest survival probability, while the 3.8 ppt and 5.7 ppt salinity treatments had lower survival than the 1.9 ppt salinity treatment (**Figure 1;** For coefficients and pairwise comparisons see supplement **Table S2**).

**Figure 1.**
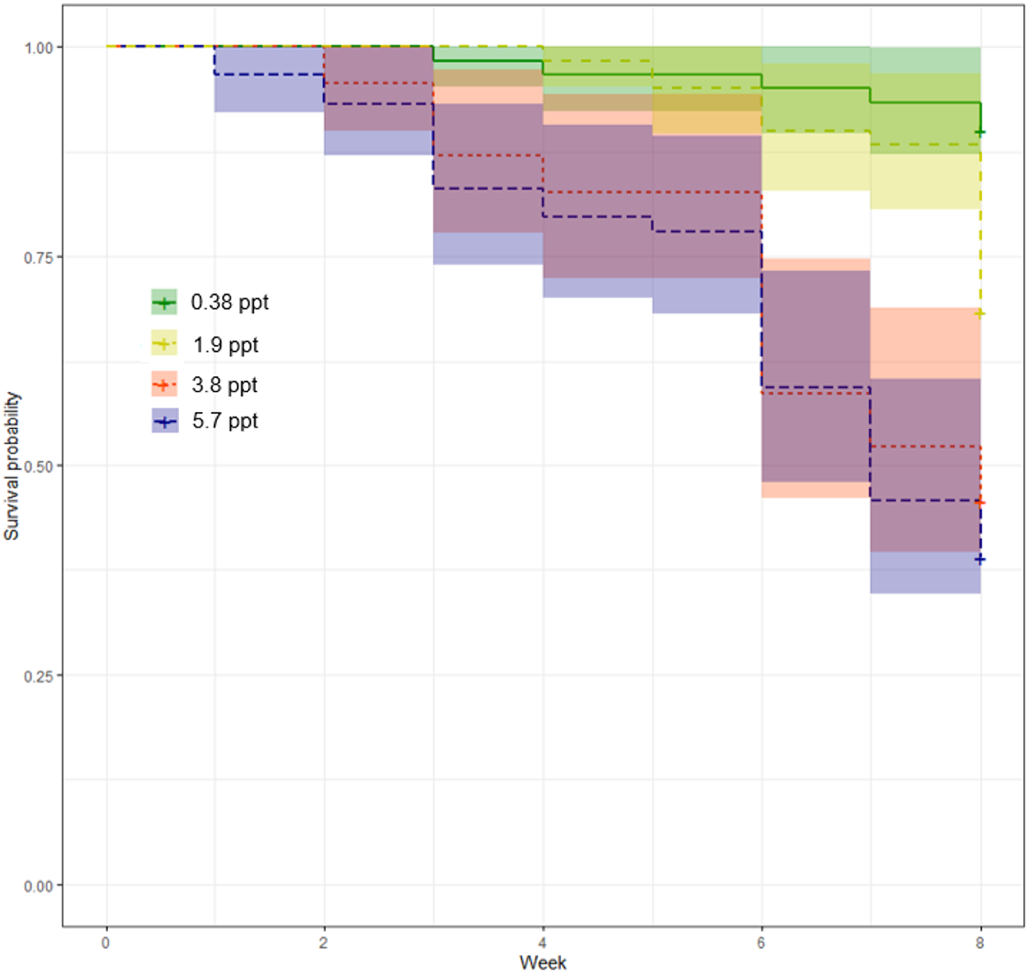
Probability of survival for snails in treatment groups of 0.38 ppt, 1.9 ppt, 3.8 ppt, 5.7 ppt salinity treatments. Snail survival in 0.38 ppt is significantly higher than 1.9 ppt (P < 0.05), 3.8 ppt (P < 0.0001), and 5.7 ppt (P < 0.0001). The survival probability of 1.9 ppt is significantly higher than 3.8 ppt and 5.7 ppt salinity treatments (P < 0.05). For coefficients and pairwise comparisons see supplement **Table S2**.

### Effect of seawater on snail egg production

Snails infected with *Schistosoma mansoni* eventually become castrated and cease producing egg masses [27]. However, since infection prevalence was lower in the 1.9 ppt, 3.8 ppt, and 5.7 ppt treatments, the decrease in egg mass production was likely in response to increases in salinity (**Figure 2**). Snails from the 0.38 ppt control treatment had higher egg output than snails in the 3.8 ppt (P < 0.05) and 5.7 ppt (P < 0.0001) salinity treatments. The 1.9 ppt treatment had a higher output than 5.7 ppt (P < 0.0001), and 3.8 ppt has a higher output than 5.7 ppt (P < 0.05). For coefficients and pairwise comparisons see supplement **Table S3**.

**Figure 2.**
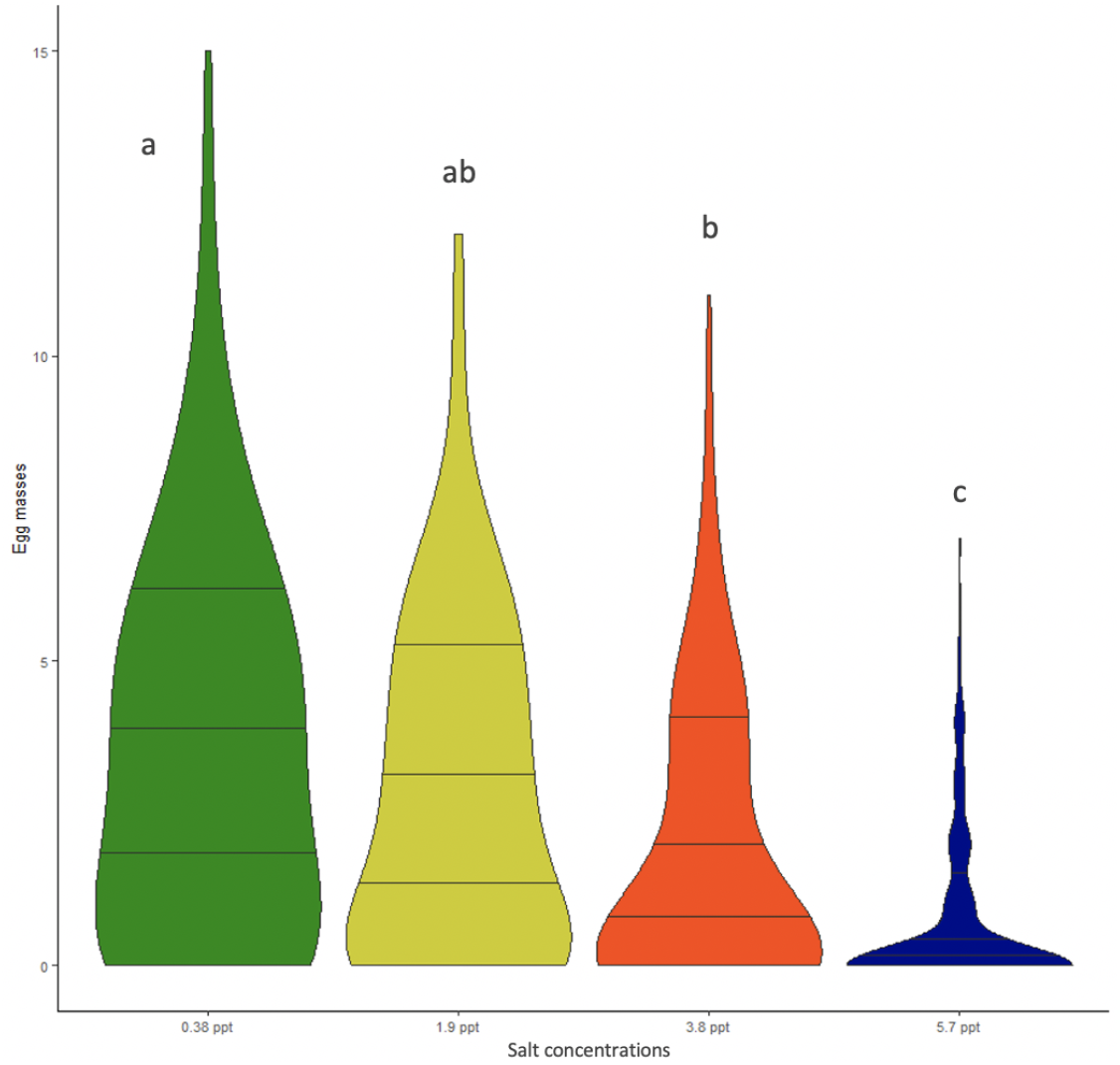
Egg mass output for uninfected snails in treatment groups 0.38 ppt, 1.9 ppt, 3.8 ppt, and 5.7 ppt. Snails in the 0.38 ppt salinity treatment had significantly higher egg mass output than 3.8 ppt (P < 0.05) or 5.7 ppt (P < 0.0001). Snails in the 1.9 ppt and 3.8 ppt treatments also had a higher output compared to the 5.7 ppt treatment (P < 0.0001 and P < 0.05, respectively). The horizontal lines on the violin plot represent data quantiles of 25%, 50%, and 75%. Letters above bars indicate significant differences in egg mass production, with different letters representing significant differences. For pairwise comparisons see supplement **Table S3**.

### Effect of seawater on infection prevalence

Infection prevalence differed among the salinity treatments with a 70% prevalence in the 0.38 ppt treatment, 16% for 1.9 ppt treatment, 13% for 3.8 ppt treatment, and 9% for 5.7 ppt treatment. Snail infection prevalence in the 0.38 ppt control treatment had significantly higher prevalence than the other 3 treatments (P < 0.0001; For coefficients and pairwise comparisons see supplement **Table S4**). However, the prevalence levels in the 1.9 ppt, 3.8 ppt, and 5.7 ppt salinity treatments were not significantly different from each other (P > 0.05) (**Figure 3**).

**Figure 3.**
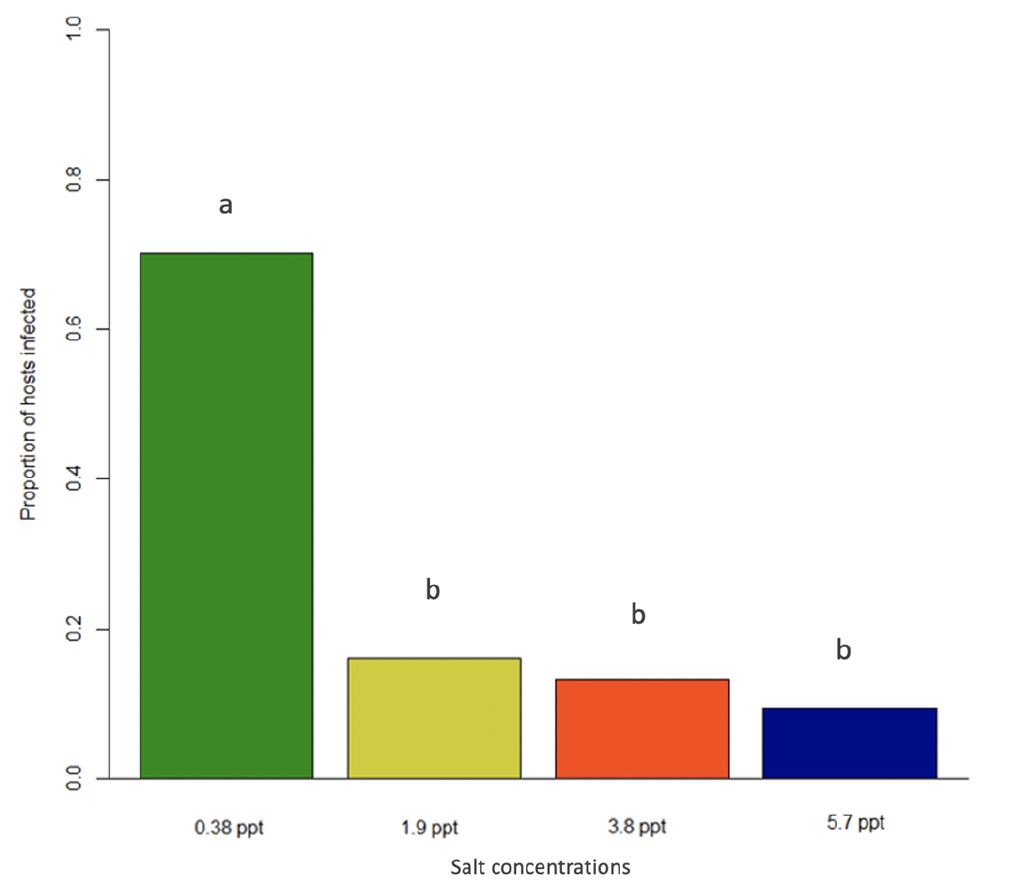
Snail infection prevalence in 0.38 ppt, 1.9 ppt, 3.8 ppt, and 5.7 ppt salinity treatments after exposure to *Schistosoma mansoni* miracidia. Different letters above bars indicate significant differences in infection prevalence among the treatments. For pairwise comparisons see supplement **Table S4**.

### Effect of seawater on cercarial survival

Cercariae in the 3.8 ppt salinity treatment had the highest survival rate. The 5.7 ppt salinity treatment had the next highest survival rate, followed by the 1.9 ppt treatment (**Figure 4**). Interestingly, the cercarial survival was lowest in the 0.38 ppt (control) treatment with this treatment having significantly lower survival than the 1.9 ppt (P < 0.05), 3.8 ppt (P < 0.0001), and 5.7 ppt treatments (P = 0.0001; For coefficients and pairwise comparisons see supplement **Table S5**). The 1.9 ppt salinity treatment was significantly lower than the 3.8 ppt treatment (P < 0.05) but was not different from the 5.7 ppt treatment (P > 0.05). The 3.8 ppt salinity treatment was significantly higher than the 5.7 ppt treatment (P < 0.05).

**Figure 4.**
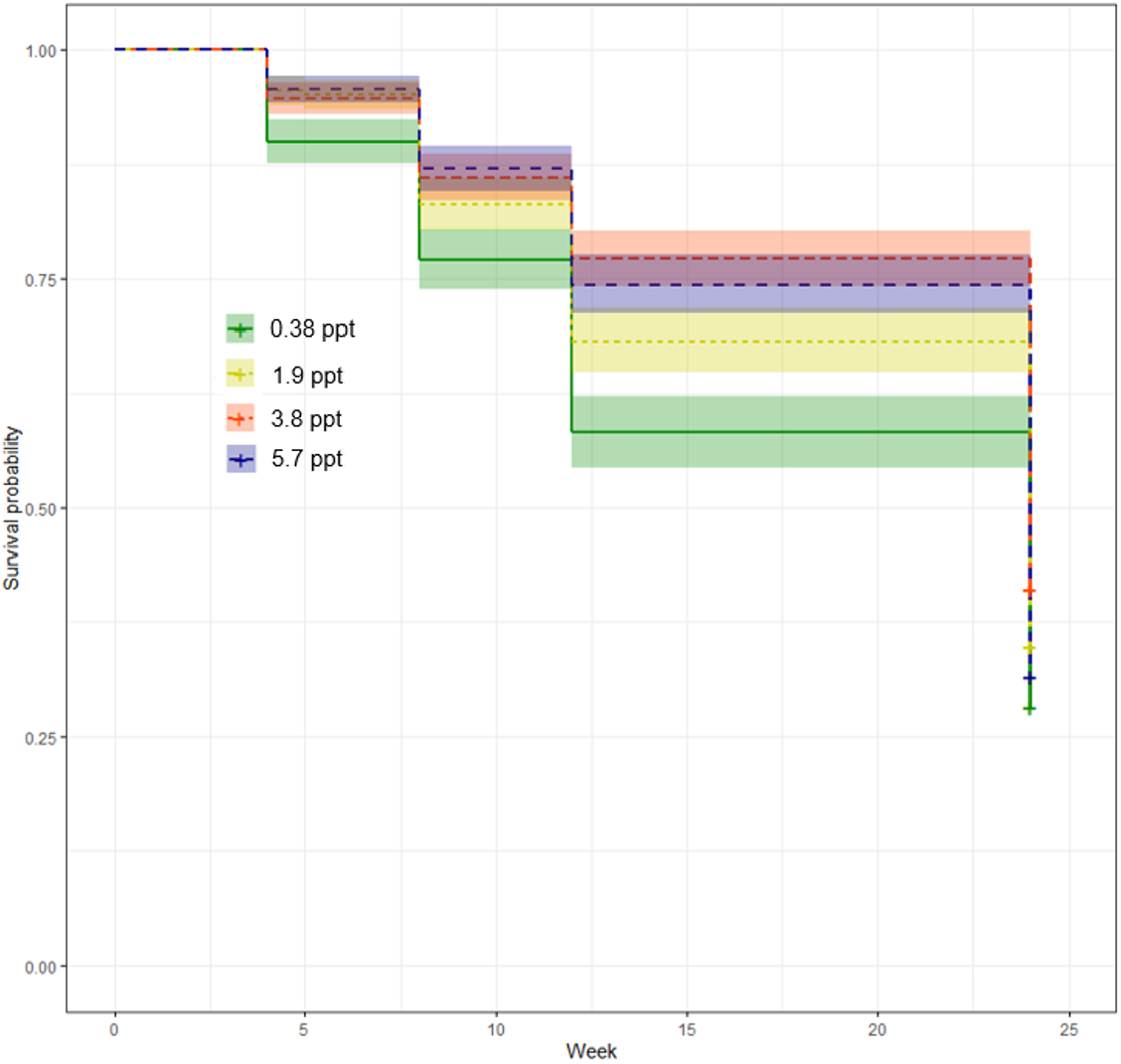
Survival probability of cercariae in 0.38 ppt, 1.9 ppt, 3.8 ppt, and 5.7 ppt salinity treatments over a 24-hour period. All four treatment groups are significantly different from one another except for 1.9 ppt and 5.7 ppt (P > 0.05) For pairwise comparisons see supplement **Table S5**.

## Discussion

Seawater intrusion caused by climate change is an on-going issue in the Nile Delta of Egypt. How seawater intrusion will impact freshwater parasite-host interactions and disease prevalence is of importance, especially in the transmission of human schistosomiasis. In this study, we explored the effect of seawater on *Schistosoma mansoni* infection success in the freshwater snail *Biomphalaria alexandrina*. Experimental conditions were designed to mimic seawater intrusion as it occurs in the Nile Delta of Egypt. We investigated crucial factors that contribute to host and parasite interactions such as snail survival, snail egg mass production, infection prevalence, as well as parasite (cercarial) survival. Our results demonstrate that snail survival (**Figure 1)**, snail reproduction (**Figure 2**), and snail infection prevalence (**Figure 3**) decreased as seawater concentrations increased across treatment groups. Additionally, cercarial survival showed a nonlinear response to seawater concentrations with the 0.38 ppt treatment having the lowest survival while cercariae in 3.8 ppt had higher survival than those in 0.38, 1.9, and 5.7 ppt treatments (**Figure 4**).

To our knowledge, this study is the first to investigate the impact of seawater on the *Biomphalaria alexandrina* – *Schistosoma mansoni* host-parasite interaction using environmentally realistic seawater solutions [28]. Previous studies have shown that the fecundity and survival of snails is adversely affected by salinities as low as 1 ppt, with significant reductions occurring between 3.5 and 4.5 ppt resulting in progressive elimination of snails [14,29]. Additionally, our result demonstrating higher cercarial survival rates at intermediate seawater concentrations is supported by previous studies, however, the salinity at which survival peaks varies among different host and parasite species and strains [30–32]. Despite the observed decreases in snail fitness and parasite transmission at higher seawater concentrations, production of infective cercariae proceeded successfully in concentrations of seawater up to 5.7 ppt suggesting that although parasite burden may be lessened, infections could still occur as sea levels rise [15 and the current study]. Of course, cercarial survival is not necessarily predictive of cercarial infectivity. This limits our ability to fully quantify the impact of rising sea levels may have on parasite transmission to humans.

We speculate that these results are caused primarily by osmotic stress and energy allocation to osmoregulation. Osmoregulation is a critical, energy-costly function of a normal cell to maintain fitness [33]. Organisms under osmotic pressure will have less energy to allocate to growth and reproduction. In our experiment, this likely led to the decrease in snail survival and snail egg mass production. Parasite larvae, such as miracidia that infect snails, are also affected by osmotic stress in the process of finding and infecting hosts, causing lower infection prevalence as salinity increased. Some authors have suggested that in contrast to miracidia, cercariae, which infect humans, require less energy for osmoregulation due to the lower difference between external and internal salinity to a certain threshold [34]. Therefore, cercariae seem to possess a higher tolerance than their snail hosts to increasing salinity, which may drive the non-linear relationship between cercarial survival and seawater concentration [34 and references therein].

## Conclusions

Taken together, reduced survival, reproduction, and infection prevalence in snail hosts with increasing salinity will lead to lower snail population sizes and potentially fewer schistosomiasis infections in areas with seawater intrusion. However, the ability of organisms to rapidly adapt to changing conditions cannot be overlooked. Although a decrease in human schistosomiasis might be expected, higher salt concentrations did not completely halt snail reproduction, snail infection, or cercarial release, suggesting parasites could still potentially infect humans, continue the life cycle, and adapt to alterations in salinity.

We have shown that increasing concentrations of seawater in freshwater systems (such as those which will occur with rising sea levels) can have a significant impact on host-parasite interactions. Here, we have focused solely on the interactions between host and parasite, but these interactions are nested within complex food webs. Parasites and their hosts can also function as prey in an ecosystem, and the changing salinity can affect these relationships and parasite transmission [36,37]. Certainly, additional factors should be evaluated to accurately assess the future trend of schistosomiasis transmission in areas with rising sea levels. Besides salinity, climate change will also influence temperature, pH, rainfall, flooding, and drought [35]. Snail fitness is likely impacted by temperature alterations and drought, and these alterations certainly impact snail mortality and parasite production [20,38–43]. Our study reveals how a single abiotic factor, salinity, can play a significant role in disease transmission. Further investigation of the role of multiple environmental factors, food web interactions, and rapid evolutionary responses of hosts and parasites to sea level rise will be needed to more accurately evaluate the future of disease transmission in these altered ecosystems.

## Acknowledgements

We thank Phillip T. LoVerde for providing the snail and parasite strain. Annabell Davis and Bailey Pyle provided invaluable assistance with data collections. Thanks to the Minchella lab members, Annabell Davis, Bailey Pyle, Hannah Melchiorre, Grace Schumacher, Spencer Siddons, and Paradyse Blackwood for reading early drafts of this manuscript.

This research did not receive any specific grant from funding agencies in the public, commercial, or not-for-profit sectors.

## Notes

### Competing Interest Statement

The authors have declared no competing interest.

## Reference

1. Zickfeld K, Solomon S, Gilford DM. Centuries of thermal sea-level rise due to anthropogenic emissions of short-lived greenhouse gases. Proc. Natl. Acad. Sci. U. S. A. 2017;114: 657–662. doi:10.1073/pnas.1612066114

2. Kaushal SS, Groffman PM, Likens GE, Belt KT, Stack WP, Kelly VR, et al. Increased salinization of fresh water in the Northeastern United States. Proc. Natl. Acad. Sci. U. S. A. 2005;102: 13517–13520. doi:10.1073/pnas.0506414102

3. Lei F, Poulin R. Effects of salinity on multiplication and transmission of an intertidal trematode parasite. Mar. Biol. 2011;158: 995–1003. doi:10.1007/s00227-011-1625-7

4. Preston D, Johnson P. Ecological consequences of parasitism. In: Nature Education Knowledge. 2010; 1:39. Available: https://www.nature.com/scitable/knowledge/library/ecological-consequences-of-parasitism-13255694/

5. Laidemitt MR, Anderson LC, Wearing HJ, Mutuku MW, Mkoji GM, Loker ES. Antagonism between parasites within snail hosts impacts the transmission of human schistosomiasis. Elife. 2019;8. doi:10.7554/eLife.50095

6. Frihy OE. The nile delta-Alexandria coast: Vulnerability to sea-level rise, consequences and adaptation. Mitig Adapt Strateg Glob Chang. 2003;8: 115–138. doi:10.1023/A:1026015824714

7. Sefelnasr A, Sherif M. Impacts of seawater rise on seawater intrusion in the Nile Delta aquifer, Egypt. Groundwater. 2013;52: 264–276. doi:10.1111/gwat.12058

8. Caffrey CR. Schistosomiasis and its treatment. Future Med Chem. 2015;7: 675– 676. doi:10.4155/fmc.15.27

9. Elmorshedy H, Bergquist R, El-Ela NEA, Eassa SM, Elsakka EE, Barakat R. Can human schistosomiasis mansoni control be sustained in high-risk transmission foci in Egypt? Parasites and Vectors. 2015;8: 372. doi:10.1186/s13071-015-0983-2

10. Barakat RMR. Epidemiology of schistosomiasis in Egypt: travel through time: review. J Adv Res. 2013;4: 425–432. doi:10.1016/j.jare.2012.07.003

11. El-Rawy M, Abdalla F, El Alfy M. Water resources in Egypt. Springer, Cham; 2020. pp. 687–711. doi:10.1007/978-3-030-15265-9_18

12. Kefford BJ, Nugegoda D. No evidence for a critical salinity threshold for growth and reproduction in the freshwater snail Physa acuta. Environ Pollut. 2005;134: 377–383. doi:10.1016/j.envpol.2004.09.018

13. Madsen H. The effect of sodium chloride concentration on centration on growth and egg laying of Helisoma duryi, Biomphalaria alexandrina and Bulinus truncatus (Gastropoda Planorbidae). J Molluscan Stud. 1990;56: 181–187. doi:10.1093/mollus/56.2.181

14. Donnelly FA, Appleton CC, Schutte CHJ. The influence of salinity on the ova and miracidia of three species of Schistosoma. Int J Parasitol. 1984;14: 113–120. doi:10.1016/0020-7519(84)90037-7

15. Chernin E, Bower C. Experimental transmission of schistosoma mansoni in brackish waters. Parasitology. 1971;63: 31–36. doi:10.1017/S0031182000067378

16. Mei B, Zhou S. Effects of osmolarity and pH on the transformation of Schistosoma japonicum miracidium to mother sporocyst in vitro. 1988. Available: http://en.cnki.com.cn/Article_en/CJFDTotal-HBYK198803001.htm

17. Seppälä O, Liljeroos K, Karvonen A, Jokela J. Host condition as a constraint for parasite reproduction. Oikos. 2008;117: 749–753. doi:10.1111/j.0030-1299.2008.16396.x

18. Kumlu M, Eroldogan OT, Aktas M. Effects of temperature and salinity on larval growth, survival and development of Penaeus semisulcatus. Aquaculture. 2000;188: 167–173. doi:10.1016/S0044-8486(00)00330-6

19. Tucker MS, Karunaratne LB, Lewis FA, Freitas TC, Liang Y san. Schistosomiasis. Curr Protoc Immunol. 2013;103. doi:10.1002/0471142735.im1901s103

20. Gleichsner AM, Cleveland JA, Minchella DJ. One stimulus-two responses: host and parasite life-history variation in response to environmental stress. Evolution (N Y). 2016;70: 2640–2646. doi:10.1111/evo.13061

21. Kassambara A, Kosinski M, Biecek P. survminer:drawing survival curves using “ggplot2”. R package version 0.4.8. J. 9, 378–400. 2020. Available: https://cran.r-project.org/package=survminer

22. Therneau TM, Lumley T, Elizabeth A, Cynthia C. Survival analysis in R. Modern Clinical Trial Analysis. 2020. doi:10.1007/978-1-4614-4322-3_1

23. Brooks, M.E., Kristensen, K., van Benthem, K.J., Magnusson, A., Berg, C.W. N A., Skaug, H.J., Maechler, M., Bolker BM. glmmTMB balances speed and flexibility among packages for zero-inflated generalized linear mixed modeling. R J. 9, 378–400. 2017.

24. Therneau TM. coxme: mixed effects cox models. 2020. Available: https://cran.r-project.org/web/packages/coxme/index.html. Version 2.2-16.

25. Lenth R, Buerkner P, Herve M, Love J, Riebl H, Singmann H. emmeans: estimated marginal means, aka least-squares means. Aka Least-Squares Means. 2020. p. https://cran.r-project.org/package=emmeans. doi:10.1080/00031305.1980.10483031>.Version1.5.0.

26. R Core Team. R: A language and environment for statistical computing. R Foundation for Statistical Computing, Vienna, Austria. 2020. Available: https://www.r-project.org/.

27. Sorensen RE, Minchella DJ. Snail-trematode life history interactions: Past trends and future directions. Parasitology. Cambridge University Press; 2001. pp. S3– S18. doi:10.1017/s0031182001007843

28. Buss N, Nelson KN, Hua J, Relyea RA. Effects of different roadway deicing salts on host-parasite interactions: The importance of salt type. Environ Pollut. 2020;266. doi:10.1016/j.envpol.2020.115244

29. Donnelly FA, Appleton CC, Schutte CHJ. The influence of salinity on certain aspects of the biology of Bulinus (Physopsis) africanus. Int J Parasitol. 1983;13: 539–545. doi:10.1016/S0020-7519(83)80025-3

30. Milotic D, Milotic M, Koprivnikar J. Effects of road salt on a free-living trematode infectious stage. J Helminthol. 2020. doi:10.1017/S0022149X20000309

31. Asch HL. Effect of selected chemical agents on longevity and infectivity of Schistosoma mansoni cercariae. Exp Parasitol. 1975;38: 208–216. doi:10.1016/0014-4894(75)90023-5

32. Venable DL, Gaudé AP, Klerks PL. Control of the Trematode Bolbophorus confusus in Channel Catfish Ictalurus punctatus Ponds Using Salinity Manipulation and Polyculture with Black Carp Mylopharyngodon piceus. J World Aquac Soc. 2000;31: 158–166. doi:10.1111/j.1749-7345.2000.tb00349.x

33. Finan JD, Guilak F. The effects of osmotic stress on the structure and function of the cell nucleus. Journal of Cellular Biochemistry. NIH Public Access; 2010. pp. 460–467. doi:10.1002/jcb.22437

34. Donnelly FA, Appleton CC, Schutte CHJ. The influence of salinity on the cercariae of three species of Schistosoma. Int J Parasitol. 1984;14: 13–21. doi:10.1016/0020-7519(84)90005-5

35. Adekiya TA, Aruleba RT, Oyinloye BE, Okosun KO, Kappo AP. The effect of climate change and the snail-schistosome cycle in transmission and bio-control of schistosomiasis in Sub-Saharan Africa. Int J Environ Res Public Health. 2020;17: 181. doi:10.3390/ijerph17010181

36. Johnson PTJ, Dobson A, Lafferty KD, Marcogliese DJ, Memmott J, Orlofske SA, et al. When parasites become prey: Ecological and epidemiological significance of eating parasites. Trends Ecol Evol. 2010;25: 362–371. doi:10.1016/j.tree.2010.01.005

37. Sokolow SH, Huttinger E, Jouanard N, Hsieh MH, Lafferty KD, Kuris AM, et al. Reduced transmission of human schistosomiasis after restoration of a native river prawn that preys on the snail intermediate host. Proc Natl Acad Sci U S A. 2015;112: 9650–9655. doi:10.1073/pnas.1502651112

38. Paull SH, Johnson PTJ. High temperature enhances host pathology in a snail-trematode system: possible consequences of climate change for the emergence of disease. Freshw Biol. 2011;56: 767–778. doi:10.1111/j.1365-2427.2010.02547.x

39. Studer A, Thieltges D, Poulin R. Parasites and global warming: net effects of temperature on an intertidal host–parasite system. Mar Ecol Prog Ser. 2010;415: 11–22. doi:10.3354/meps08742

40. Poulin R. Global warming and temperature-mediated increases in cercarial emergence in trematode parasites. Parasitology. 2006;132: 143–151. doi:10.1017/S0031182005008693

41. McCreesh N, Booth M. The effect of simulating different intermediate host snail species on the link between water temperature and schistosomiasis risk. Munderloh UG, editor. PLoS One. 2014;9: e87892. doi:10.1371/journal.pone.0087892

42. Mas-Coma S, Valero MA, Bargues MD. Climate change effects on trematodiases, with emphasis on zoonotic fascioliasis and schistosomiasis. Vet Parasitol. 2009;163: 264–280. doi:10.1016/j.vetpar.2009.03.024

43. Mangal TD, Paterson S, Fenton A. Predicting the impact of long-term temperature changes on the epidemiology and control of schistosomiasis: a mechanistic model. Baylis M, editor. PLoS One. 2008;3: e1438. doi:10.1371/journal.pone.0001438

